# Assessing PDB Macromolecular Crystal Structure Confidence at the Individual Amino Acid Residue Level

**DOI:** 10.1101/2022.05.17.492280

**Authors:** Chenghua Shao, Sijian Wang, Stephen K. Burley

## Abstract

Approximately 87% of the more than 190,000 atomic-level, (three-dimensional) 3D biostructures in the Protein Data Bank (PDB) were determined using macromolecular crystallography (MX). Agreement between 3D atomic coordinates and experimental data for >100 million individual amino acid residues occurring within ∼150,000 PDB MX structures was analyzed in detail. The Real-Space-Correlation-Coefficient (RSCC) calculated using the 3D atomic coordinates for each residue and experimental electron density enables outlier detection of unreliable atomic coordinates (particularly important for poorly-resolved sidechain atoms) and ready evaluation of local structure quality by PDB users. For human protein MX structures in PDB, comparisons of per-residue RSCC experimental-agreement metric with AlphaFold2 computed structure model confidence (pLDDT-predicted local distance difference test) document (i) that RSCC values and pLDDT scores are correlated (median correlation coefficient∼0.41), and (ii) that experimentally-determined MX structures (3.5 Å resolution or better) are more reliable than AlphaFold2 computed structure models and should be used preferentially whenever possible.

## Introduction

The PDB was established in 1971 as the first digital data resource in biology (Protein Data Bank, 1971). It has grown more than 27,000-fold to become the only freely accessible global archive of 3D structures of proteins, nucleic acids, and their complexes with one another and small-molecule ligands experimentally determined using macromolecular crystallography (MX), nuclear magnetic resonance (NMR) spectroscopy, and electron microscopy (3DEM). Open access to well-validated, expertly-biocurated PDB structures enables scientific advances across fundamental biology, biomedicine, energy sciences, and biotechnology/bioengineering (Burley et al., 2018; Goodsell et al., 2020; Westbrook and Burley, 2019; Westbrook et al., 2020). PDB structures also played critical roles in efforts aimed at predicting (or computing) atomic-level 3D structure models from protein sequence alone (Burley and Berman, 2021; Burley et al., 2021). Today, AlphaFold2 (Jumper et al., 2021; Tunyasuvunakool et al., 2021) and RoseTTAFold (Baek et al., 2021) support computation of structure models of globular proteins with accuracies comparable to those of lower-resolution experimental methods.

The Worldwide Protein Data Bank (wwPDB, wwpdb.org; Berman et al., 2003; wwPDB consortium, 2019) manages the PDB archive according to the FACT [Fairness-Accuracy-Confidentiality-Transparency (van der Aalst et al., 2017)] and FAIR [Findable-Accessible-Interoperable-Reusable (Wilkinson et al., 2016)] principles. Current wwPDB members include RCSB Protein Data Bank or RCSB PDB (Berman et al., 2000; Burley et al., 2022; Rose et al., 2021); Protein Data Bank in Europe (Mir et al., 2018); and Protein Data Bank Japan (Kinjo et al., 2018), plus two specialist data resources [3DEM data resource: Electron Microscopy Data Bank (Abbott et al., 2018); NMR data resource: Biological Magnetic Resonance Bank (Ulrich et al., 2008)]. The wwPDB OneDep system for global deposition, validation, and biocuration of PDB structures (Young et al., 2017; Gore et al., 2017; Young et al., 2018; Feng et al., 2021) serves tens of thousands of structural biologists working on every inhabited continent. wwPDB validation reports generated within OneDep for every PDB structure provide comprehensive quality assessments, calculated using community-standard software tools. For MX structures, wwPDB validation reports summarize individual residue quality using the local electron density goodness-of-fit metric RSCC (Tickle, 2012; Brändén and Jones, 1990).

Relentless growth in the number of MX structures in PDB since 1971 has yielded an enormous body of open access data for biomedical research. It has also created considerable challenges for some PDB data consumers, who may encounter difficulties when discerning which part (or parts) of a given PDB structure are not to be trusted, and, consequently, may not be useful for interpreting experimental results or generating new hypotheses. With open access to hundreds of thousands of computed structure models of proteins from AlphaFoldDB (Varadi et al., 2022) and the ModelArchive (Schwede et al., 2009), choosing atomic-level 3D structure information on which to rely on has become even more challenging. Herein, we describe (1) use of RSCC to identify very well-resolved and well-resolved regions of MX structures in PDB, and (2) how RSCC compares with the AlphaFoldDB computed structure model prediction confidence metric pLDDT for all MX structures of human proteins represented in PDB.

## Results

### RSCC Distribution

#### (A) RSCC Distribution and Identification of Outliers in PDB MX Structures at the Individual Residue Level

The distribution of RSCC values for >100 million standard amino acid residues and nucleotides from ∼150,000 PDB MX structures is illustrated in Figure 1A (mean and median values 0.935 and 0.955, respectively). Markedly different from a normal distribution, the skewed RSCC distribution is heavily tailed on its left-side (lower RSCC values). Therefore, neither standard deviation (*σ*) nor interquartile range (IQR) can be used to accurately characterize statistical dispersion or identify outliers within the RSCC distribution. For example, >4% of residues in PDB have RSCC values below μ(mean)-2*σ, versus* only 2.5% for a normal distribution. An alternative criterion for outlier classification frequently used for non-normal symmetrical distributions is 1.5 IQR below the first quartile. Because the RSCC distribution is so heavily skewed, opting for this metric would classify >7% of residues as RSCC outliers. Transformation of the RSCC distribution into a regular parametric distribution was attempted unsuccessfully. By way of explanation, the Pearson correlation coefficient bounded between -1 and 1 can be transformed into normal distribution through Fisher’s Z-transformation, but only when the data used to generate the correlation coefficient have a bivariate normal distribution. This condition is not met for either experimentally-observed electron density or calculated electron density based on 3D structure atomic coordinates.

**Figure 1.**
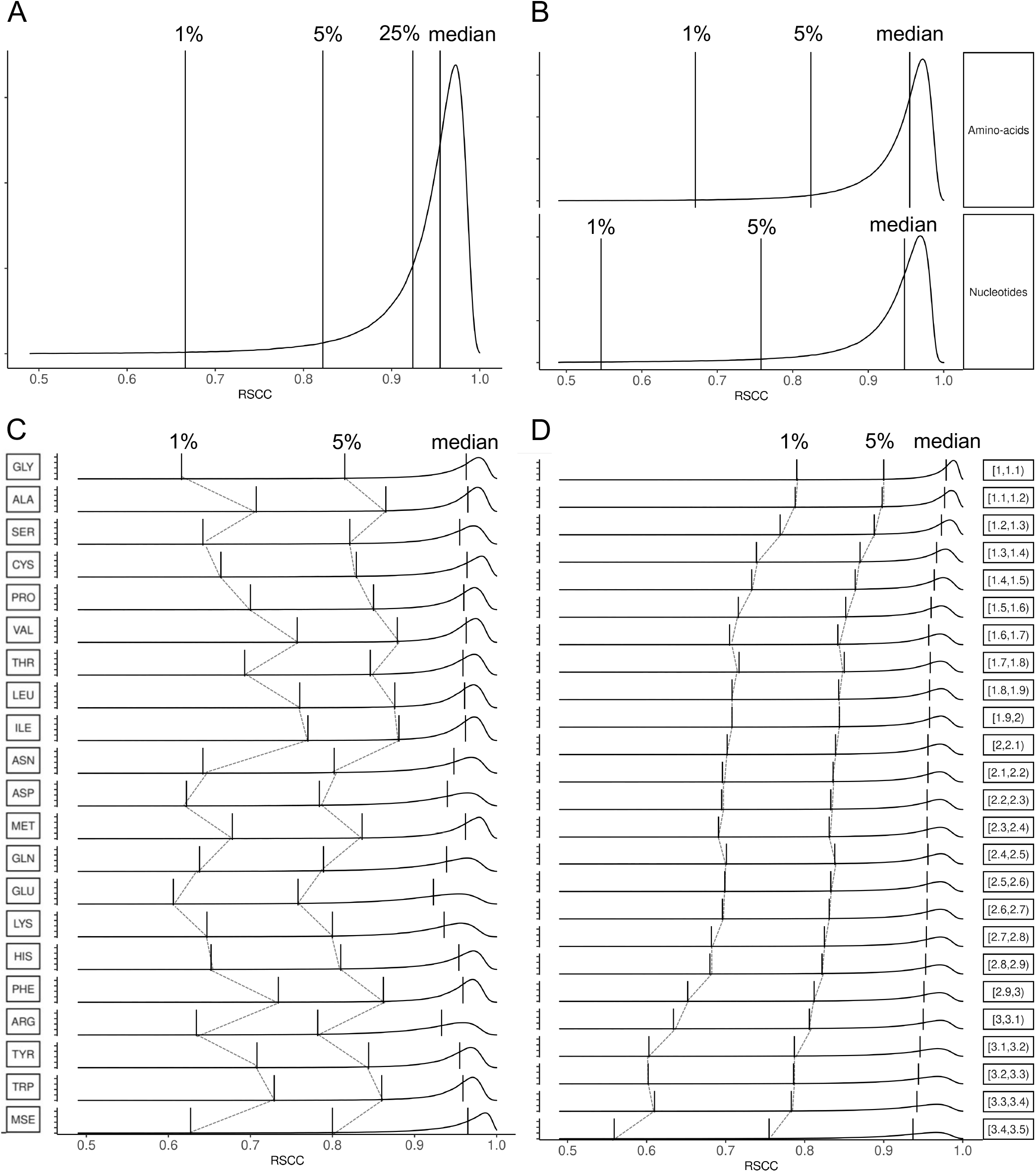
RSCC value distribution of standard residues in the PDB archive. Probability density plots of RSCC values for (A) all standard protein and nucleic acid residues, (B) amino acid residues *versus* nucleotides, (C) individual amino acid types ordered by number of atoms (plus MSE seleno-methionine), and (D) individual amino acid types *vs*. resolution limit. Vertical lines denote the median value, and the 5% and 1% percentiles, respectively (*i*.*e*., cumulative percentages of data values to the left of each line). 25% (1^st^ quartile) line is shown in (A).

An outlier is defined as an observation that deviates so much from the other observations as to arouse suspicion that it originated from a different mechanism (Hawkins, 1980). Using a probability density-based approach, outliers constitute the least probable observations associated with the lowest estimated probability density from the data distribution (Shao et al., 2018). RSCC values follow a unimodal distribution with monotonically increasing probability density from the lowest value to the mode or peak value (Figure 1A). Consequently, for data values less than the mode the ranking of the probability density is consistent with the ranking of the data itself from low to high, permitting identification of one-sided outliers with a percentile cut off. The lowest 1% of RSCC values (Figure 1A, to the left of vertical 1% line) have the lowest probability, and the lowest 5% RSCC values (Figure 1A, between 1% and 5% lines) have low probability. Both 1% and 5% probability cutoffs are commonly used thresholds for classifying data values as outliers. The remainder of the first quartile (between 5% and 25%) can be considered as having intermediate probability.

#### (B) RSCC Distribution versus Structure Residue Type and MX Resolution Limit

RSCC values depend on the chemical nature of the biopolymer component. Comparison of RSCC distributions for proteins and nucleic acids (Figure 1B) documents that RSCC values for nucleotides are typically lower than for amino acid residues, indicating that the fit of atomic coordinates to experimental electron density for individual nucleotides is generally inferior to that observed for individual amino acid residues in PDB MX structures. Figure 1C illustrates RSCC distributions for each amino acid residue type (hereafter residue) separately. In general, residues with non-polar sidechains that occur more frequently within the hydrophobic cores of globular proteins (*e*.*g*., valine, leucine, isoleucine, phenylalanine, and tryptophan) have higher median RSCC values than residues with polar sidechains more frequently found on the surfaces of globular proteins (*e*.*g*., serine, asparagine, aspartate, glutamine, glutamate, lysine, histidine, and arginine). The relative paucity of steric constraints on surface residue atomic positions permits adoption of multiple sidechain and even backbone conformations that may not be well resolved in the MX experiment, particularly at resolution limits lower than ∼3.5 Å. Analyses presented below were limited to PDB MX structures of proteins.

Resolution limit of the experimental diffraction data represents an important determinant of MX structure quality, both globally and locally. Figure 1D illustrates RSCC distributions *versus* resolution limit. As expected, the median value of RSCC decreases with resolution limit, because experimental electron density is less well resolved at lower resolution. This trend is most evident when comparing higher- (1.0-1.5 Å) and lower-resolution limit ranges (3.0-3.5 Å). RSCC distributions for each residue type were evaluated *versus* resolution limit (see Supplementary Data spreadsheet for tabulated summary statistics). These results enable identification of lowest and low probability RSCC value outliers for each residue type as a function of resolution limit. [N.B.: Unlike small-molecule crystallography, a high-precision/high-accuracy experimental method, very few PDB MX structures (∼0.5%) were determined at resolution limits better than 1.0 Å.]

For the avoidance of doubt, MX is an extremely powerful experimental tool for resolving 3D structures of well-ordered, globular proteins. Residues with RSCC values greater than 0.85 can be trusted for any residue type. The fraction of all individual residues in PDB MX structures with RSCC>0.85 is ∼95% for those with resolution limits better than 2 Å (∼50% of PDB MX structures) and ∼93% for those with resolution limits better than 3.5 Å (∼98% of PDB MX structures), documenting the high resolving power of the method. In contrast, MX is not well suited to the challenge of resolving conformational heterogeneity within proteins, whatever its origins. The atomic coordinates of poorly-resolved individual residues present in a given PDB MX structure (*i*.*e*., outlier residues with lowest and low probability RSCC values) should not trusted.

### Comparing RSCC Distributions with AlphaFoldDB pLDDT Distributions

#### (A) Comparing RSCC and pLDDT for PDB MX Structures of Human Proteins

To assess 3D structure prediction confidence quantitatively, AlphaFold2 provides per-residue pLDDT scores (scaled between 0 and 100): very high confidence if pLDDT≥90; confident if 90>pLDDT≥70; low confidence if 70>pLDDT≥50; very low confidence if pLDDT<50. Artificial intelligence/deep learning approaches have also been shown to outperform physicochemical based methods for predicting intrinsically-disordered regions (IDRs) of proteins (Necci et al., 2021). Among them, lower pLDDT scores have been shown to be a good predictors of protein disorder (Ruff and Pappu, 2021).

More than 23,000 AlphaFold2 predicted 3D structures of human proteins (computed structure models or CSMs) were downloaded from AlphaFoldDB for analysis. The pLDDT distribution of ∼15 million individual residues contained within the downloaded CSMs is illustrated Figure 2 (dashed line). It is bimodal (major peak∼95, minor peak∼35). Approximately 28% of downloaded CSM residues have pLDDT<50, indicating very low confidence in their predicted atomic coordinates, which is consistent with earlier observations (Thornton et al., 2021) and independent estimates of IDRs in the human proteome (Ruff and Pappu, 2021; Tunyasuvunakool et al., 2021).

**Figure 2.**
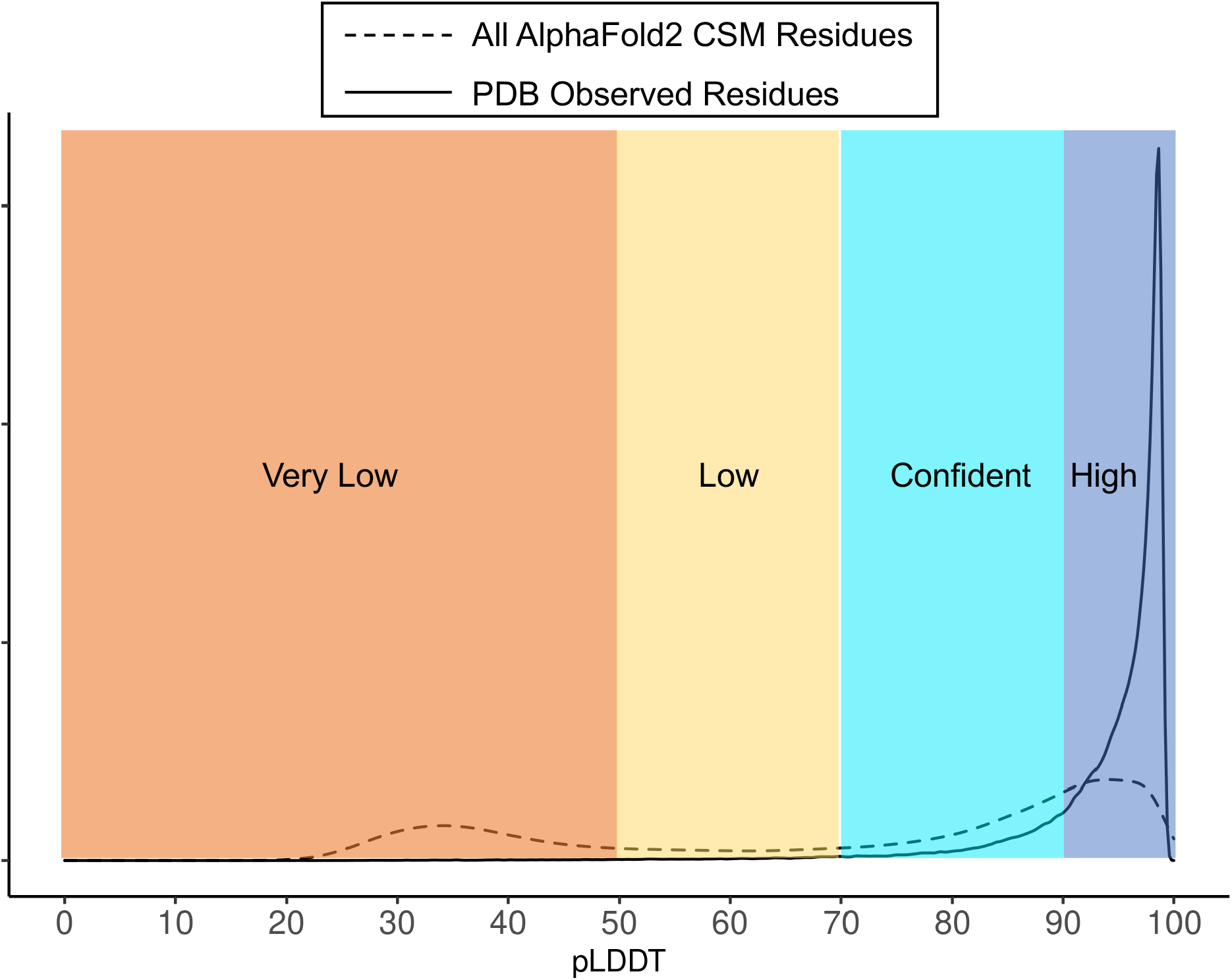
pLDDT score distributions of AlphaFoldDB CSMs of human proteins. Dashed Line: Human protein residues (major peak∼95, minor peak∼35). Solid: Human protein residues observed in PDB MX structures. Prediction confidence color coding: blue-very high confidence pLDDT≥90; cyan-confidence 70≤pLDDT<90; yellow-low confidence 50≤pLDDT<70; orange-very low confidence pLDDT<50.

Amino acid sequences of each human protein AlphaFoldDB CSM were used to query and align with all PDB protein structure sequences using the RCSB PDB 1D coordinate server Application Programming Interface or API (https://1d-coordinates.rcsb.org/) (Segura et al., 2020). Approximately 7,500 unique human protein sequences were detected in more than 53,000 PDB structures. A majority of human protein structures in PDB encompass only individual domains of longer polypeptide chains (∼64% of structures have <95% sequence coverage), with the remainder spanning the entire polypeptide chain (∼36% of structures have ≥95% sequence coverage).

To compare RSCC and pLDDT at the individual residue level, RSCC values for complete residues (*i*.*e*., excluding all residues with missing atoms, partial atomic occupancy, and/or multiple conformations) in over 41,000 PDB MX structures of ∼5,300 unique human proteins were selected and compared to pLDDT scores of the corresponding AlphaFoldDB CSMs in pairwise fashion. The pLDDT distribution for complete residues occurring in PDB MX structures is unimodal (solid line in Figure 2; median pLDDT score∼96, ∼2.4% residues have pLDDT<70, ∼0.6% residues have pLDDT<50). These findings will not surprise MX practitioners. The method performs best with crystalline samples of relatively compact globular proteins (lacking poorly ordered N-or C-termini and/or long surface loops). Considerable efforts in expression construct design are frequently required before MX can be employed for high-resolution structural studies of human protein domain structures (Gao et al., 2005).

#### (B) Comparing RSCC and pLDDT for Full-length Human RNA-Binding Protein Nova-1

Overall correlation coefficient between per residue RSCC values and pLDDT scores (RSCC/pLDDT-CC) for every human protein MX structure in PDB is plotted in Supplementary Figure S1 (median value∼0.41, range -0.48 to 0.95). RSCC and pLDDT were compared for representative individual proteins on a per residue basis. Figure 3 illustrates our findings for full-length human RNA-binding protein Nova-1 (UniProt ID P51513), which consists of an N-terminal segment plus three globular K-homology or KH domains (KH1, KH2, and KH3), well separated from one another in amino acid sequence (Figure 3A). Two related MX structures are available from the PDB [PDB: 2ANR (KH1 and KH2) (Teplova et al., 2011); and PDB: 1DT4 (KH3) (Lewis et al., 1999)]. To compare RSCC and pLDDT graphically, pLDDT was scaled by 1/100 (resulting metrics falling between 0 and 1) and per residue values were plotted as a function of UniProt sequence numbering (Figure 3B).

**Figure 3.**
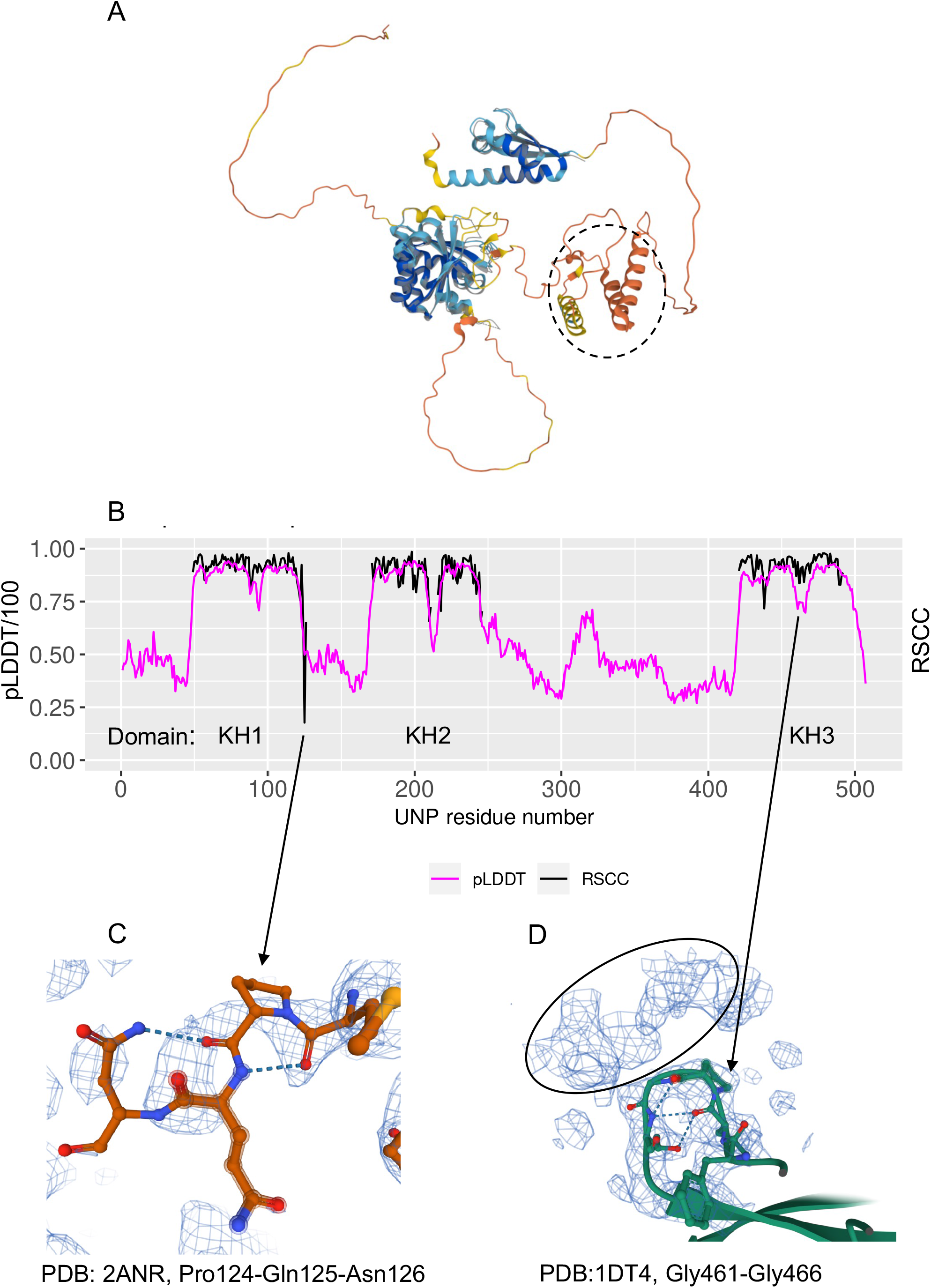
Human RNA-binding protein Nova-1: Comparison of RSCC values for two PDB structures and pLDDT scores of the AlphaFoldDB CSM. (A) Mol* (Sehnal et al., 2021) graphical display of the AlphaFoldDB CSM of the complete polypeptide chain, color coded by pLDDT scores as for Figure 2. PDB: 2ANR (KH1 and KH2) and PDB: 1DTJ (KH3) superposed on the CSM as half-transparent gray shade. Dashed line circle denotes a portion of the full-length human Nova-1 protein not previously recognized as potentially globular. (B) Per-residue overlay of pLDDT/100 (magenta, full-length UniProt ID P51513) and RSCC (black, observed residues in PDB: 2ANR (KH1 and KH2) and PDB: 1DT4 (KH3)]. The gap in the KH2 overlay corresponds to an unobserved polypeptide chain segment in PDB: 2ANR. (C) Experimental electron density overlayed on PDB: 2ANR atomic coordinates for residues 124-126 of domain KH1 (2|F_observed_|-|F_calculated_| map, contoured at 1.0 *σ*). (D) Experimental electron density overlayed on PDB: 1DT4 (KH3) atomic coordinates for loop residues 461-466 (2|F_observed_|-|F_calculated_| map, contoured at 1.0 *σ*), which are stabilized by the crystal contacts with a neighboring protomer (see electron density within black ellipse).

Figure 3B shows that per residue RSCC values for PDB: 2ANR (KH1 and KH2) and pLDDT scores for the same residues in the AlphaFoldDB CSM are well correlated (RSCC/pLDDT-CC∼0.75 for common residues). Close inspection of the experimental electron density for KH1 and KH2 domain residues revealed that most were well resolved (*i*.*e*., high RSCC values). Minor exceptions include the loops connecting the second and third *β*-strands in the three-stranded, anti-parallel *β*-sheet characteristic of KH domains (Lewis et al., 1999), and the C-terminus of KH1 (Figure 3C). Limited proteolysis studies of full-length human Nova-1 protein documented susceptibility to cleavage in these same regions (see Figure 2 of (Lewis et al., 1999)), suggesting that they are conformationally flexible in solution. The AlphaFoldDB CSM for human Nova-1 superposes well on PDB: 2ANR (KH1 and KH2) with C*α* Root-Mean-Square-Difference or RMSD∼0.3 Å. Residues in the inter-strand loops and the limited number of residues inter-domain region that were observed in PDB: 2NAR (KH1 and KH2), albeit with low RSCC values, and predicted in the AlphaFoldDB CSM with low pLDDT scores exhibit somewhat different conformations, providing further evidence that these segments of the human Nova-1 polypeptide chain are flexible.

Figure 3B also reveals that per residue RSCC for PDB: 1DT4 (KH3) and pLDDT scores for the same residues in the AlphaFoldDB CSM are not as well correlated as seen for domains KH1 and KH2 (RSCC/pLDDT-CC∼0.39). The AlphaFoldDB CSM and PDB: 1DT4 (KH3) superpose well (C*α* RMSD∼0.7 Å). Structural differences between PDB: 1DT4 (KH3) and the AlphaFoldDB CSM are restricted to the inter-strand loop (residues 461-466), the position of which appears to be stabilized by interactions with a neighboring protomer in the crystal lattice (Figure 3D). Again, these findings suggest that KH domain inter-strand loops are conformationally flexible.

#### (C) Comparing RSCC and pLDDT for Individual Human Proteins

RSCC values for all complete residues in PDB MX structures of human proteins are correlated with pLDDT scores for the same residues in the AlphaFoldDB CSMs (RSCC/pLDDT-CC∼0.30) (Supplementary Data spreadsheet). Close examination of additional representative PDB MX human structures and their AlphaFoldDB CSMs confirms that polypeptide chain segments with RSCC<0.8 and pLDDT<50 typically correspond to internal loop regions or N- and C-termini of the structure. Residues with higher RSCC values (>0.9) generally correspond to residues with higher pLDDT scores (>70) for most PDB MX structures of human proteins, except for polypeptide chain segments that participate in intermolecular interactions with other proteins or nucleic acids, crystal packing interactions, or contacts with small-molecule ligands. Backbone atoms in AlphaFoldDB CSMs with median pLDDT>90 generally superpose well on their corresponding PDB MX structures, even in cases when the RSCC/pLDDT-CC is lower (*i*.*e*., 0.2-0.4).

Table 1 summarizes results obtained by comparing the AlphaFoldDB CSM for human hemoglobin subunit alpha (HbA-*α*, UniProt ID P69905; median of individual residue pLDDT scores∼98.6) and PDB MX structures of the same protein in various oxidation states. Three sets of atomic coordinates for HbA-*α* were extracted from PDB MX structures of the hemoglobin *α*_2_*β*_2_ hetero-tetramer in the following oxidation states: PDB: 2DN1 (oxy); PDB: 2DN2 (deoxy); and PDB: 2DN3 (carbonmonoxy). All three structures were experimentally determined at ∼1.25 Å resolution by the same research group with well-resolved electron density for most non-hydrogen atoms and median RSCC values>0.94 (Park et al., 2006). The chain backbone of the single AlphaFoldDB CSM for HbA-*α* superposes well on those of all three experimental structures (C*α* RMSD∼0.3 to ∼0.5 Å) with RSCC/pLDDT-CC ranging from ∼0.2 to ∼0.6. RSMD values between each of the three PDB structures and the AlphaFoldDB CSM calculated using non-hydrogen atoms are higher than those obtained for C*α* atoms alone (ranging from ∼0.7 to ∼1.5 Å). These results document that predicted sidechain atomic positions in the AlphaFoldDB CSM for HbA-*α* are less accurate than those of main chain atoms. Supplementary Figure S2 shows that His88 (responsible for binding to the heme group iron atom) was only predicted to occur in the same position as that observed by MX in PDB: 2DN2 (deoxy), which differs in the other two oxidation state structures. In the same view, the predicted conformation of Leu83 resembles that of PDB: 2DN1 (oxy), but not the other two oxidation states. Also in the same view, the predicted conformation of Trp14 differs dramatically from that observed in PDB: 2DN1 (oxy). Therefore, even AlphaFoldDB CSMs with very high median pLDDT scores cannot be relied upon to reproduce the “ground-truth” of well-determined, high-resolution PDB MX structures, particularly when the macromolecules in question are structurally dynamic.

**Table 1:** Comparison of the AlphaFoldDB CSM and three high-resolution PDB structures of human HbA-*α*.

| PDB MX Structure | Resolution Limit | Oxidation State | Median RSCC | Overall RSCC/pLDDT-CC | RMSD ($C\alpha$ Å) | RMSD (Non-H Å) |
| --- | --- | --- | --- | --- | --- | --- |
| 2DN1 | 1.25 Å | oxy | 0.97 | 0.59 | 0.53 | 1.45 |
| 2DN2 | 1.25 Å | deoxy | 0.94 | 0.16 | 0.29 | 0.71 |
| 2DN3 | 1.25 Å | carbonmonoxy | 0.97 | 0.57 | 0.54 | 1.32 |

#### (D) Contrasting RSCC and pLDDT for Individual Human Proteins

Most PDB human protein backbone structures superpose well on their corresponding AlphaFoldDB CSMs, and RSCC values are generally correlated with pLDDT scores. However, discordance between RSCC values and pLDDT scores does occur. We analyzed human proteins for which AlphaFoldDB CSMs exhibited very low pLDDT scores while corresponding PDB MX structures (containing the same polypeptide chain or segment thereof) had high RSCC values, and *vice versa*.

First, AlphaFoldDB CSMs for human proteins with low overall pLDDT scores were examined and compared to their PDB MX structure counterparts. 71 PDB MX structures have corresponding AlphaFoldDB CSMs with median pLDDT<50 (Supplementary Data spreadsheet), which differ significantly in 3D from one another. For example, PDB: 3LK3 (Hernandez-Valladares et al., 2010), determined at 2.68 Å resolution, encompasses residues 971-1035 of UniProt ID Q5VZK9 (human F-actin-uncapping Leucine-rich repeat-containing protein 16A; entity ID 3; median RSCC∼0.95). The corresponding AlphaFoldDB CSM has very low confidence (pLDDT<50 for every residue; median pLDDT∼36). The AlphaFoldDB CSM and the PDB MX structure are too different to be superposed in 3D. The AlphaFoldDB CSM prediction shows only random coil (Supplementary Figure S3). PDB: 3LK3 is a multi-protein, co-crystal structure of this polypeptide chain segment of UniProt ID Q5VZK9 bound to its target (F-actin-capping protein). In the co-crystal, two shorter polypeptide chain segments (971-1004 and 1021-1035) were experimentally observed as a U-shaped, mostly random coil sitting in a groove formed by approximation of two of its known target proteins (Supplementary Figure S3). The corresponding experimental electron density is well resolved, except for two residues (Cys1004 and Arg1021), both of which have RSCC<0.8 and flank residues 1005-1020 that were not experimentally observed.

Low pLDDT score segments within AlphaFoldDB CSMs do not always correspond to intrinsically disordered regions (IDRs) of proteins. The PDB MX structure of human PR domain zinc finger protein 4 (UniProt ID Q9UKN5 residues 393-530; PDB: 3DB5; DOI: 10.2210/pdb3DB5/pdb), determined at 2.15 Å resolution (median RSCC∼0.96), reveals a compact, largely *β*-strand domain (Supplementary Figure S4). The corresponding AlphaFoldDB CSM has median pLDDT∼37, indicating very low confidence in the prediction, which does include some *β*-strand-like features, albeit with a different topological arrangement in 3D from PDB: 3DB5 (Supplementary Figure S4). Not surprisingly, therefore, the C*α* RMSD between PDB: 3DB5 and its corresponding CSM is very high (∼17 Å). PDB: 3DB5 is a high-quality structure with atomic coordinates of most of its residues well resolved in the experimental electron density. Within the domain structure, residues in the *β*-strands have higher RSCC values *versus* those occurring within inter-strand loops, exemplifying the value of RSCC as a quantitative measure of experimental structure local quality. Alternatively, some AlphaFoldDB CSMs with low pLDDT scores can partially be partially superposed on their corresponding PDB MX structures, as exemplified in Supplementary Figure S5 for the CSM of UniProt ID P41182 and PDB: 7LWE.

In addition to AlphaFoldDB CSMs with very low overall pLDDT scores, individual residues with pLDDT<50 in AlphaFoldDB CSMs were examined. Particular attention was paid on those residues with pLDDT<50 present in 288 very high-resolution (1 to 1.1 Å) PDB MX structures of human proteins. Among these PDB MX structures, we identified 70 “discordant” residues with RSCC>0.95 and pLDDT<50. For example, PDB: 4FKA (Prugovecki et al., 2012) is an MX structure of human insulin determined at 1.08 Å resolution (UniProt ID P01308), within which 12 residues are well resolved in the experimental electron density but do not superpose well onto their counterparts in the AlphaFoldDB CSM. It was previously observed for this protein that “the AlphaFold model bears no resemblance to the PDB structure, possibly because it has missed the disulfide bonds that hold the protein together.” (Thornton et al., 2021). The lesson from the insulin example is the importance of preferentially using well-determined PDB structures whenever they are available, and not relying on AlphaFoldDB as a sole source of 3D structural information.

The obverse scenario of high pLDDT *versus* low RSCC was also examined for representative cases. PDB: 7E5M (Sun et al., 2021) is MX structure of residues 33-268 of human tumor-associated calcium signal transducer 2 determined at 3.2 Å resolution (UniProt ID P09758), with median RSCC∼0.67 *versus* median pLDDT∼94 for the corresponding AlphaFoldDB CSM. The experimental structure and the CSM are very similar, with C*α* RMSD∼0.5 Å when loop residues 82-103 are excluded from the 3D comparison (Supplementary Figure S6). Close inspection of the experimental electron density revealed that the lower RSCC value likely stems from the paucity of diffraction data. At resolution limits worse than 3.5 Å (*i*.*e*., higher number), MX structures are not always well resolved, because the number of experimental observations (diffraction measurements) per atom may be insufficient for the method to accurately determine atomic positions.

### RSCC-based Confidence Criterion and Color Scheme for PDB MX Structure Display

Atomic coordinates of most PDB MX structures are well resolved in the experimental electron density. Within individual MX structures, however, atomic coordinates for individual residues or short segments of the polypeptide chain(s) may not be as accurate. Statistically rigorous outlier detection of RSCC values can provide readily interpretable measures of local structure quality for PDB data consumers who are not experts in structural biology. Similar to the AlphaFoldDB pLDDT display color scheme, residues in PDB MX structures can be assigned an RSCC-based confidence and colored-coded so that RSCC outliers are readily apparent in ribbon representation 3D graphical displays. The RCSB PDB has adopted a color-coding scheme comparable to that used by AlphaFoldDB. The vast majority of residues that were very well resolved by the MX method (very well resolved - RSCC ordinal ranking between 25% and 100%, *i*.*e*., the most probable RSCC range) are colored blue. Well resolved residues with RSCC ordinal ranking between 5% and 25% are color coded cyan. Outlier residues that are not well resolved by the MX method are color coded either yellow (low confidence, RSCC ordinal ranking between 1% and 5%) or orange colored (very low confidence, lowest 1% of RSCC values).

Figure 4 illustrates application of this RSCC probabilities-based color-coding scheme for PDB: 1DTJ [third domain (KH3) of human RNA-binding protein Nova-2 determined at 2.0 Å resolution, UniProt ID Q9UNW9 (Lewis et al., 1999)]. Most human Nova-2 KH3 residues are colored blue or cyan both in 1D (Figure 4A) and 3D (Figure 4B) representations of the MX structure, reflecting the fact that they were well resolved or very well resolved by the method. Some residues occurring in the inter-strand loop, a short segment between the first and second *α*-helices, and the C-terminus of the domain are colored yellow or orange, because their RSCC values were deemed to be statistical outliers (falling within the lowest 1-5% or lowest 1% of the probability distributions for those particular amino acids at 2.0 Å resolution). The similarity of the RCSB PDB color scheme with that of Alphafold2 pLDDT confidence scores, buttressed by the comparability of the distributions depicted in Figures 1 and 2, is intended to help users of PDB data make informed assessments of experimentally-determined structure quality without having to delve into the details of the wwPDB validation report. It is also intended to aid direct comparison of PDB MX structures with CSMs of proteins generated *via* deep learning methods.

**Figure 4.**
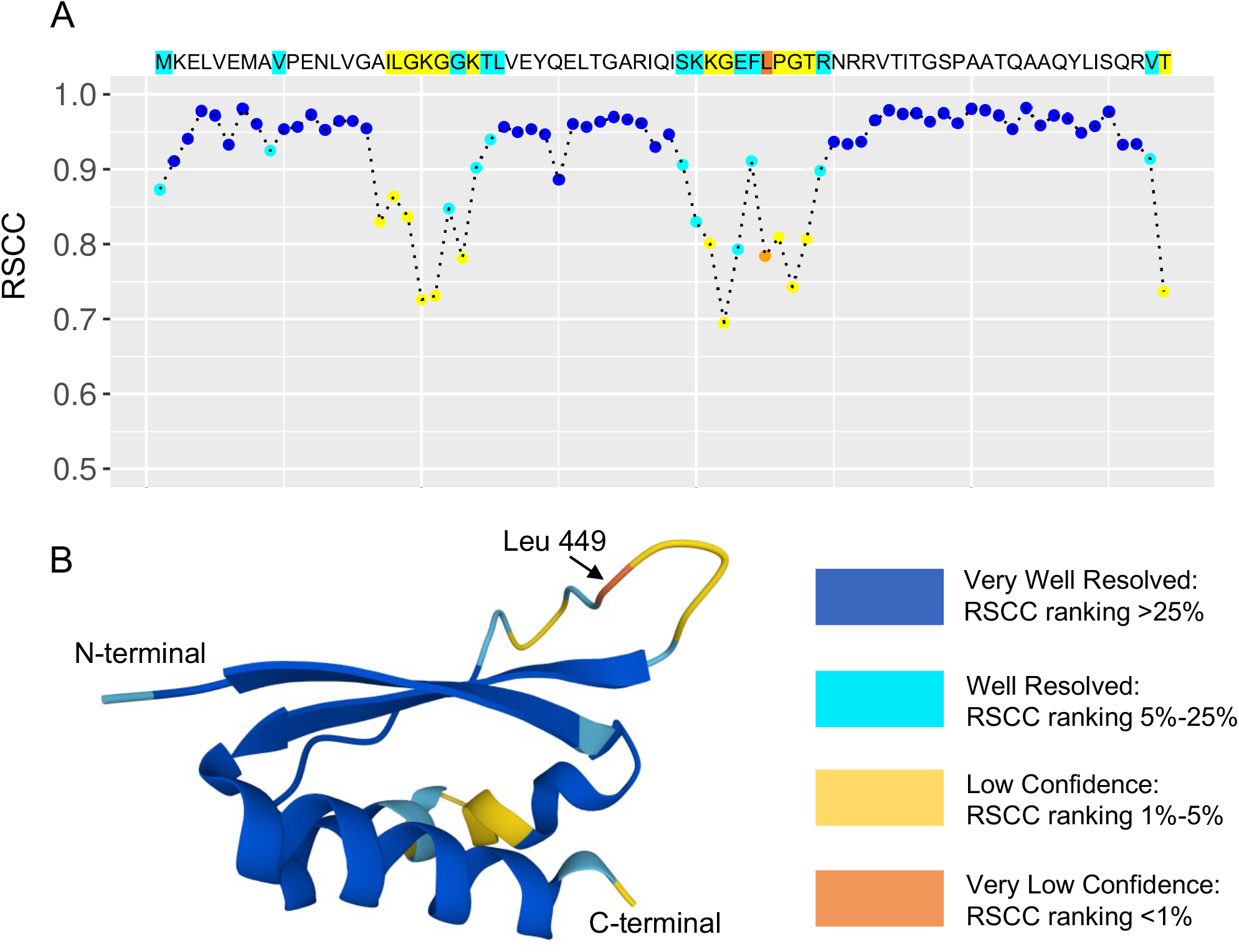
Human RNA-binding protein Nova-2 KH3 domain: RSCC-based confidence levels and corresponding color scheme for PDB MX structures. (A) Amino acid sequence of PDB: 1DTJ (KH3) serving as X-axis, each residue is denoted by a solid circle in the 2D graph based on RSCC value (Y-axis). For each residue, both its 1-letter amino acid code and circle are color coded by per-residue structure quality using RSCC probability distributions of same residue type at similar resolution limits in PDB MX structures: very well resolved (RSCC ranking >25%, high probability, blue); well resolved (RSCC ranking 5%-25%, intermediate probability, cyan); low confidence (RSCC ranking 1%-5%, low probability, yellow); very low confidence (RSCC ranking <1%, lowest probability, orange). (B) Mol* ribbon drawing of PDB: 1DTJ (KH3) using the same color scheme.

## Discussion

Analyses of the distribution of RSCC values for PDB MX structures demonstrated the utility of RSCC as a residue-level MX structure quality indicator that can be used to assess the goodness of fit of atomic coordinates to experimental electron density. Outlier residues are objectively identified as those with lowest 1% or 5% probability ranking. Such outliers typically have little, if any, corresponding experimental electron density signal. Low confidence and very low confidence portions of PDB MX structures should be treated with caution. In many, perhaps most, cases, low RSCC values indicate that the corresponding residue(s) or short polypeptide chain segments are not well ordered in the crystal, and do not, therefore, contribute to the Bragg diffraction signal measured in the MX experiment. They may be statically or dynamically (*i*.*e*., flexible) disordered. The MX method cannot distinguish these possibilities.

A potentially important outcome of this work is inclusion of 3D display of RSCC outliers for PDB MX structures (*e*.*g*., Figure 4B) on the RCSB PDB research-focused web portal. This telegraphic approach to displaying structure quality information is aimed primarily at the estimated 99% of RCSB.org users who are not experts in structural biology. The blue-cyan-yellow-orange color-coding scheme based on RSCC was chosen because it should be familiar to users of CSMs generated by AlphaFold2, RoseTTAFold (Baek et al., 2021), *etc*. Residues that were very well resolved or well resolved by the MX experiment (identified in blue and cyan, respectively) represent ∼95% of those found in PDB MX structures with resolution limits better than 3.5 Å, reflecting the high overall quality of 3D structure data archived in the PDB. The remaining 5% of residues are color coded yellow for low confidence (low probability RSCC ordinal ranking of 1%-5%) or orange for very low confidence (lowest probability RSCC ordinal ranking up to 1%). The atomic coordinates of these poorly resolved residues are not to be trusted. Many structural biologists are taking a “glass half full” view of CSMs generated using AlphaFold2, RoseTTAFold, *etc*. They use the CSMs of full-length eukaryotic proteins to design protein expression constructs that exclude low confidence (50≤pLDDT<70, color coded yellow) and very low confidence (pLDDT<50, color coded orange) segments of longer polypeptide chains to generate samples of truncated proteins well suited for structure/function studies using MX, NMR, or 3DEM. They also scrutinize segments of polypeptide chains with low and/or very low confidence predictions for potentially globular segments that have not been previously characterized (within the dashed line circle in Figure 3A, for example). For the other 99% of PDB data consumers, poorly resolved residues in PDB MX structures can be viewed in two ways. They can be seen as an inconvenience because they are not reliable. Alternatively, they can be viewed as a source of opportunities for designing experiments using methods other than MX to probe the biological or biochemical function of poorly resolved residues.

To explore the question of disorder in MX structures, RSCC values were compared and contrasted to the AlphaFoldDB CSM pLDDT scores for all human protein structures in the PDB, considering both entire structures and individual residues. RSCC values in PDB structures and AlphaFoldDB CSM pLDDT scores for the same human proteins are correlated, suggesting that both metrics can be used to assess polypeptide chain flexibility or disorder. Cases wherein RSCC values and AlphaFoldDB CSM pLDDT scores are not correlated may serve as useful case studies for those seeking to improve *de novo* protein structure prediction methods.

## STAR Methods

### RESOURCE AVAILABILITY

#### Lead Contact

Further information and requests for resources should be directed to and will be fulfilled by the Lead Contact, Dr. Chenghua Shao.

#### Materials Availability

This study did not use or generate any physical material.

#### Data and Code Availability

- PDB structure and validation data utilized in this study are available through FTP at ftp.wwpdb.org and through HTTP at RCSB.org under individual PDB IDs.
- This paper does not report original code.
- Any additional information required to reanalyze the data reported in this paper is available from the lead contact upon request.

### EXPERIMENTAL MODEL AND SUBJECT DETAILS

All data are generated from the datasets provided in the KRT (to be submitted when accepted).

### METHOD DETAILS

#### Data Collection

All data used for this study were based on the publicly released PDB archive available at ftp.wwpdb.org. Data were extracted from both atomic coordinate files and wwPDB validation reports released before Mar 18, 2022 and then aggregated through data processing. Human protein AlphaFold2 CSMs were downloaded from AlphaFoldDB (Varadi et al., 2022) on Mar 18, 2022.

#### Sequence Alignments for AlphaFoldDB CSMs and PDB Structures

Amino acid sequences corresponding to human protein AlphaFoldDB CSMs (based on UniProt ID) were used to query and align with protein sequences represented in the PDB archive using the RCSB PDB 1D coordinate server API (https://1d-coordinates.rcsb.org/) (Segura et al., 2020). PDB structure sequences were then used for residue-level alignments to match RSCC values and pLDDT scores for the same residue at the same location in each polypeptide chain.

#### Pairing RSCC and pLDDT at Residue Level for Each Human Protein Structure

Only polypeptide chain segments >30 residues in length were included in this step because correlation coefficients calculated between RSCC values and pLDDT scores may not be reliable when matched sequence regions are too short. Sequence pairing between PDB MX structures and AlphaFoldDB CSMs of human proteins also followed additional criteria enumerated below:

1. Alignment lengths between PDB structures and CSMs must be of equal length. In case of gaps or insertions, they were not paired.
2. Pairing must be on the same residue type. In case of mutations, they were not paired.
3. Paired residues must have valid RSCC values, otherwise they were not paired.
4. Pairing was performed on the first instance of a CSM, if multiple CSMs were identified.
5. Pairing was performed on the first instance of the protein in each PDB structure if multiple instances of the protein in PDB were identified.
6. Pairing was only performed on fully-resolved residues (*i*.*e*., all atoms present in the PDB structure).
7. Pairing was only performed on residues with occupancy ≥ 0.9

#### Computation and Software

Data processing, visualization, search, tabulation, and statistical calculation were performed primarily using a combination of Python and R. Calculations were performed on in-house RCSB PDB workstations.

### QUANTIFICATION AND STATISTICAL ANALYSIS

#### Data Summary

As of Mar 18, 2022, there are 164,404 PDB MX structures. Among these MX structures, 149,757 have RSCC calculated successfully on 108,286,678 standard residues in their wwPDB validation reports. The remainder do not have experimental data or do not have standard residues (*e*.*g*., carbohydrate only structures), or failed RSCC calculation. Among these standard residues, ∼97% are standard amino acids and ∼3% are standard nucleotides. Overall statistical characteristics of RSCC values are as follows: mean=0.935, median=0.955, standard deviation=0.065, IQR=0.047, 25^th^ percentile (1^st^ quartile)=0.924, 5^th^ percentile=0.822, 1^st^ percentile=0.666.

AlphaFoldDB human protein CSMs encompass 20,504 UniProt IDs and 23,391 CSMs. Only the 1^st^ isoform of the UniProt ID is used for the alignment to PDB MX structures. 7,502 human protein UniProt IDs could be aligned to 78,088 entities in 53,714 PDB structures. During RSCC/pLDDT pairing, 5,340 UniProt IDs could be aligned to 41,306 PDB structures.

#### Data Exploration and Visualization

Preliminary data exploration was carried out by running R on the dataset collected above. Tables and figures were generated use R and standard software packages. Probability density distributions were calculated using Gaussian kernel density estimate.

#### Future Implementation and Annual Update

The RSCC-based structure quality classification and color scheme will be implemented within 3D visualization tools provided on RSCB.org. Outlier criteria will be updated annually with newly deposited structures included. Hence, outlier classification of individual residues in PDB MX structures may change slightly from year to year.

## Acknowledgements

RCSB PDB is funded by the <u>National Science Foundation</u> (DBI-1832184, PI: S.K.B.), the <u>US Department of Energy</u> (DE-SC0019749, PI: S.K.B.), and the <u>National Cancer Institute, National Institute of Allergy and Infectious Diseases</u>, and <u>National Institute of General Medical Sciences</u> of the <u>National Institutes of Health</u> under grant R01GM133198 (PI: S.K.B.).

## Author Contributions

Conceptualization: S.K.B., C.S., and S.W.; Methodology: C.S., S.W., and S.K.B.; Statistical Analysis: C.S. and S.W.; Writing: C.S., S.K.B., and S.W.; Supervision: S.K.B.; Funding Acquisition: S.K.B.;

## Supplementary Data Spreadsheet

Four data sheets are provided:

RSCC_by_residue_and_resolution: Per-residue RSCC value distribution for all standard amino acid residues in all PDB MX protein structures by residue type and by resolution limit. 1%, 5%, and 25% thresholds are the percentile values from the lowest.

RSCC_pLDDT_per_PDB: Per-PDB structure comparison between per-residue RSCC values in PDB MX human protein structure and pLDDT scores of residues in the corresponding AlphaFoldDB CSM from the same sequence. Each row corresponds one PDB structure that may have >1 corresponding UniProt IDs.

RSCC_pLDDT_by_residue: Comparison between per-residue RSCC values and pLDDT scores, grouped by residue type.

RSCC_pLDDT_by_resolution: Comparison between per-residue RSCC values and pLDDT scores, grouped by resolution limit.

