## Supplementary Figures for "Assessing PDB Macromolecular Crystal Structure Confidence at the Individual Amino Acid Residue Level"

Supplementary Figure S1: RSCC/pLDDT-CC: Correlation coefficient between RSCC in PDB MX structures and pLDDT in AlphaFold2 CSMs for human proteins.

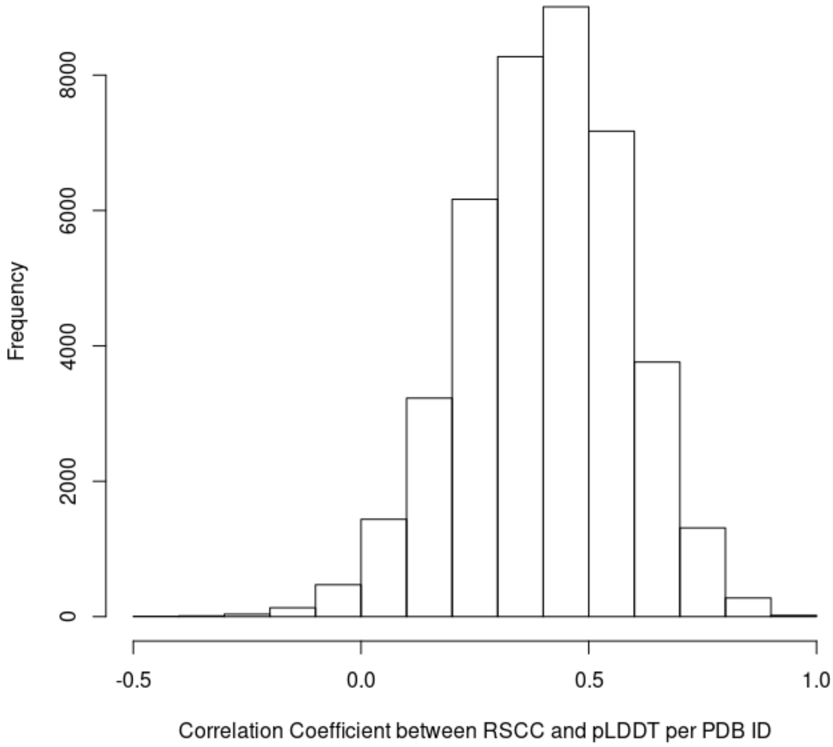

Supplementary Figure S2: Ribbon drawing overlay of human hemoglobin alpha subunit AlphaFold2 CSM (green) with PDB: 2DN1 oxy (magenta), PDB: 2DN2 deoxy (cyan), and PDB: 2DN3 carbonmonoxy (yellow). Selected sidechains represented as atomic stick figures (Carbon-ribbon color for CSM and each PDB structure; Oxygen-red; Nitrogen-blue).

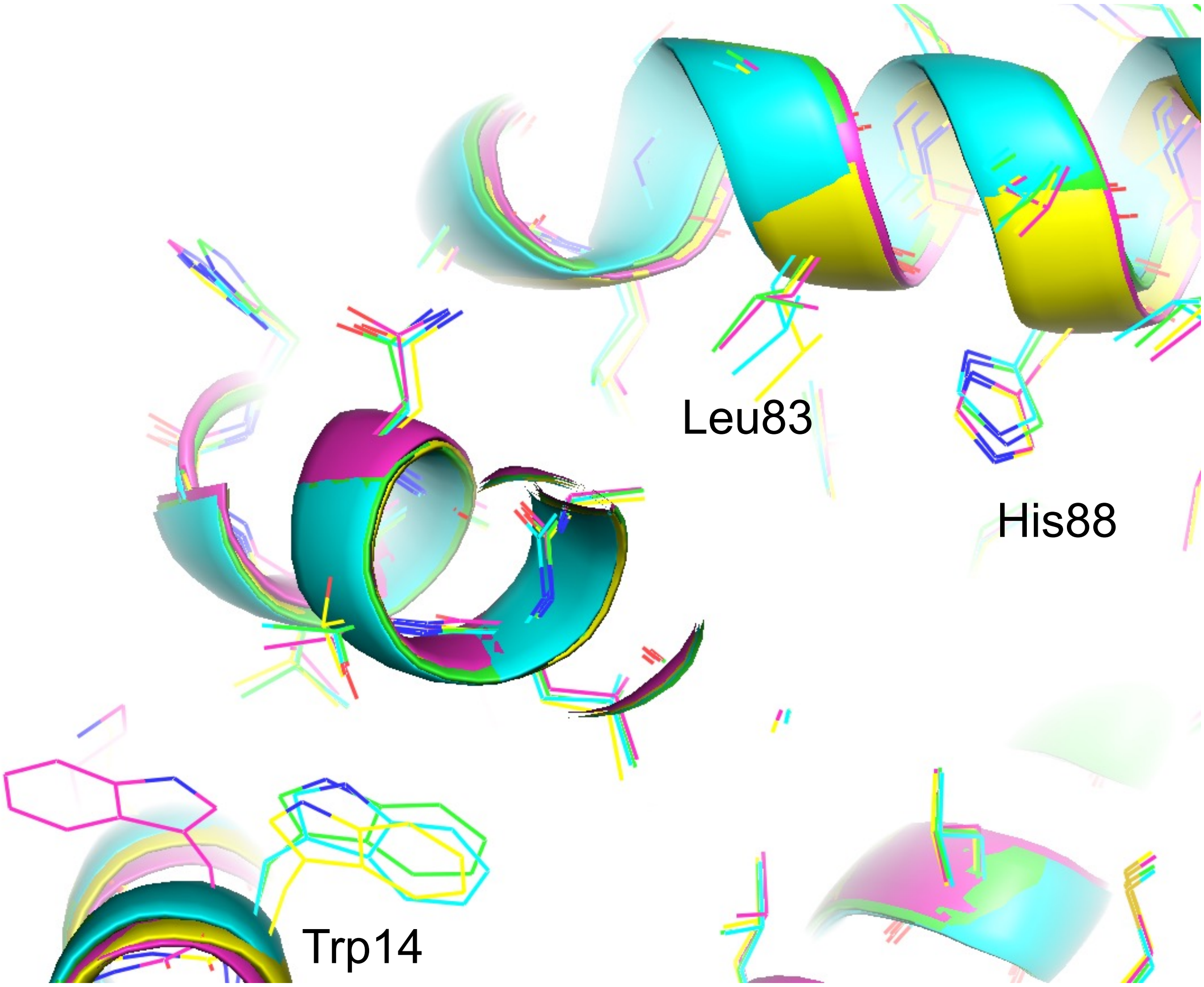

Supplementary Figure S3: Ribbon drawings of AlphaFold2 CSM for UniProt ID Q5VZK9 (left) *versus* PDB: 3LK3 (right). Structural components not shared are colored grey. CSM is colored based on pLDDT color scheme (Orange: pLDDT<50). PDB MX structure is colored green (dashed line, denotes residues not observed by MX).

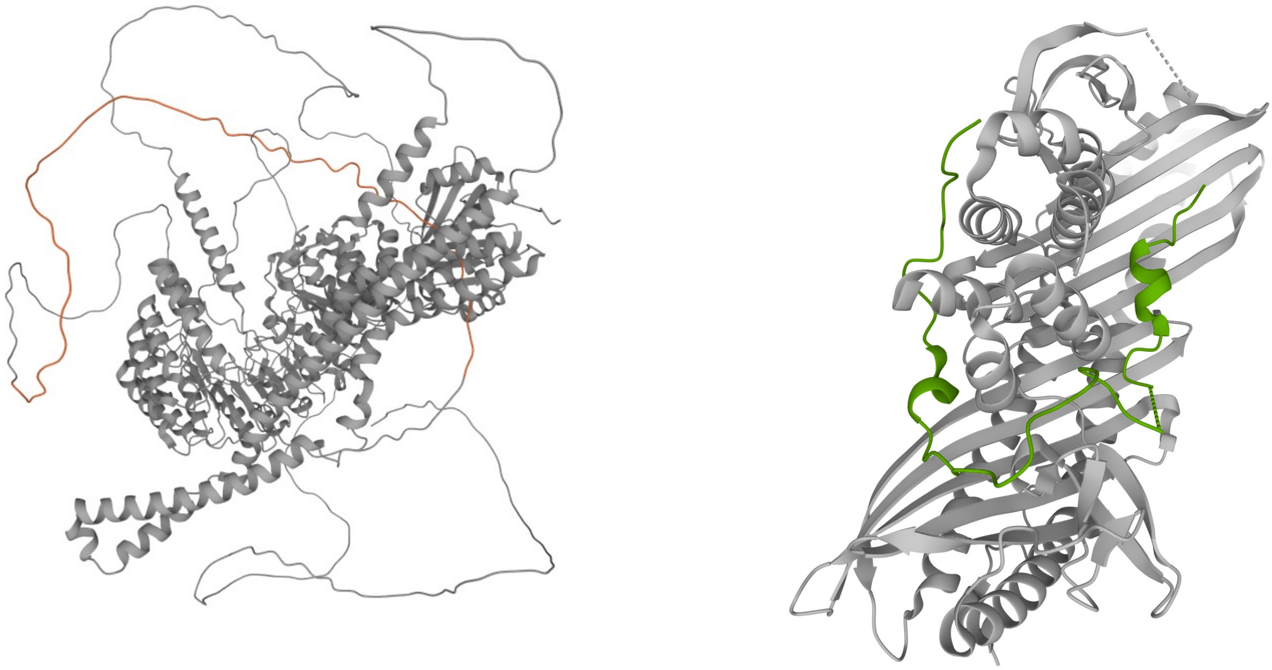

Supplementary Figure S4: Ribbon drawings of AlphaFold2 CSM for UniProt ID Q9UKN5 (left) *versus* PDB: 3DB5 (right). Structural components not shared are colored grey. CSM is colored based on pLDDT color scheme (Orange: <50; Yellow: 50 to <70). PDB MX structure is colored green (dashed line, denotes residues not observed by MX).

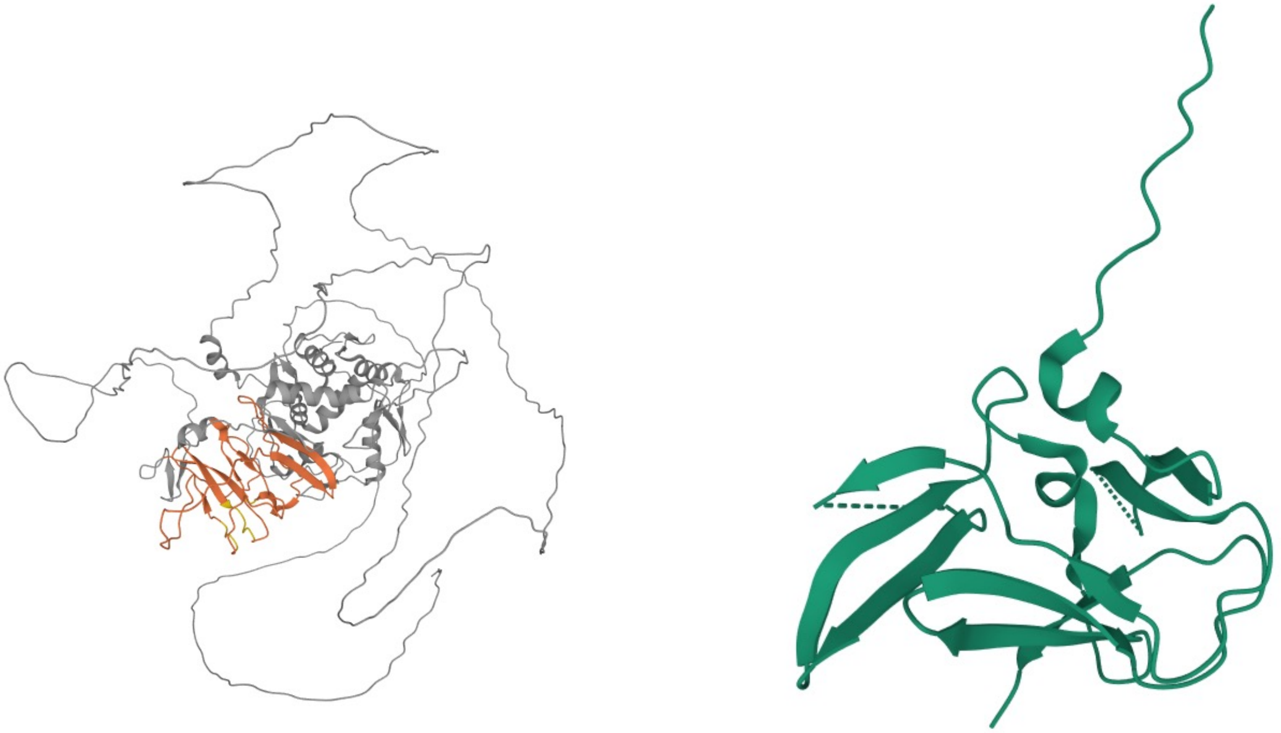

Supplementary Figure S5: Ribbon drawings of AlphaFold2 CSM on UniProt ID P41182 (left) *versus* PDB: 7LWE (right). Structural components not shared are colored grey. CSM is colored based on pLDDT color scheme (Orange: <50; Yellow: 50 to <70; Cyan: 70 to <90). PDB MX structure is colored green. The N-terminal domain of UniProt ID P4118 also known as human B-cell lymphoma 6 protein represents a useful exemplar. In all there are 67 PDB structures of this domain, which is a 2-layer sandwich of α-helices and β-sheets (Sillitoe et al., 2021). In contrast, the AlphaFold2 CSM has low confidence (median pLDDT~46; pLDDT<50 for most individual residues). Comparison of the AlphaFold2 CSM with PDB: 7LWE (determined at 1.17 Å, median RSCC~0.99) revealed that the two share an almost identical 2-helix bundle local structure for residues 101-128 (Cα RMSD~0.4 Å) but are otherwise very different in 3D (Supplementary Figure S5). The electron density for PDB: 7LWE was well resolved, except for a few N- and C-terminal residues with lower RSCC values.

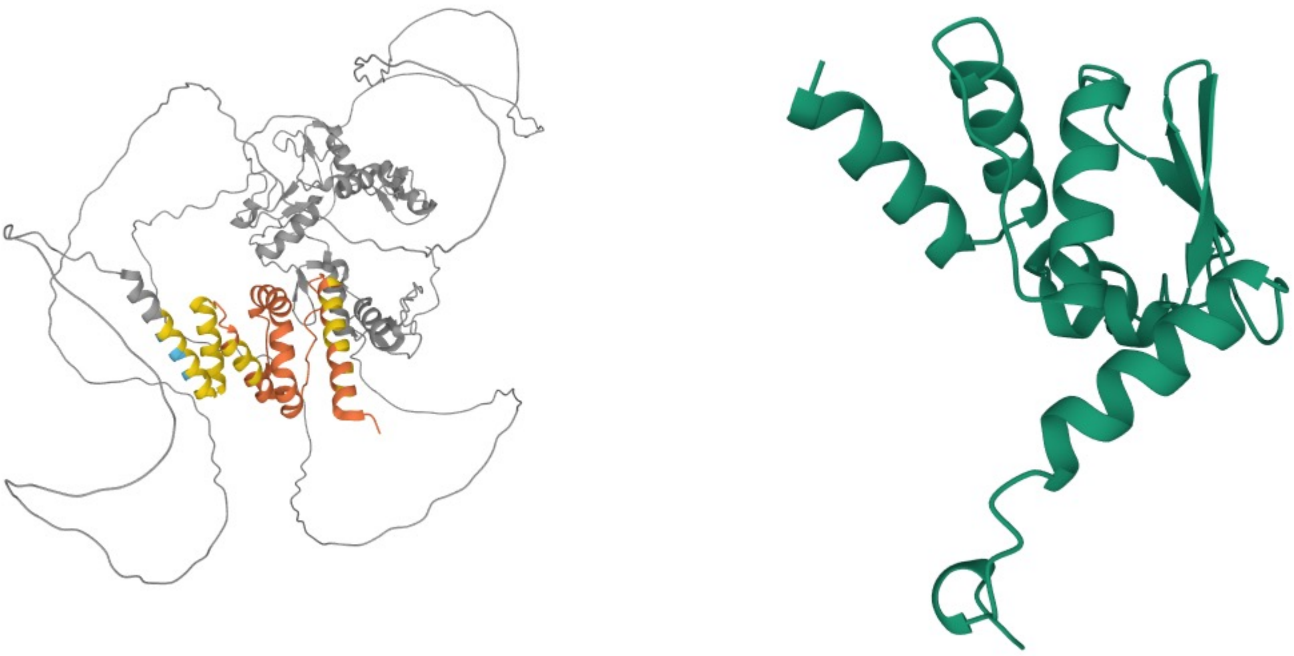

Supplementary Figure S6: Ribbon drawing overlay of AlphaFold2 CSM on UniProt ID P09758 *versus* PDB: 7E5M. Structural components not shared are colored gray. CSM is colored based on pLDDT color scheme (Orange: <50; Yellow: 50 to <70; Cyan: 70 to <90; Blue: ≥90). PDB MX structure is colored as half-transparent green.

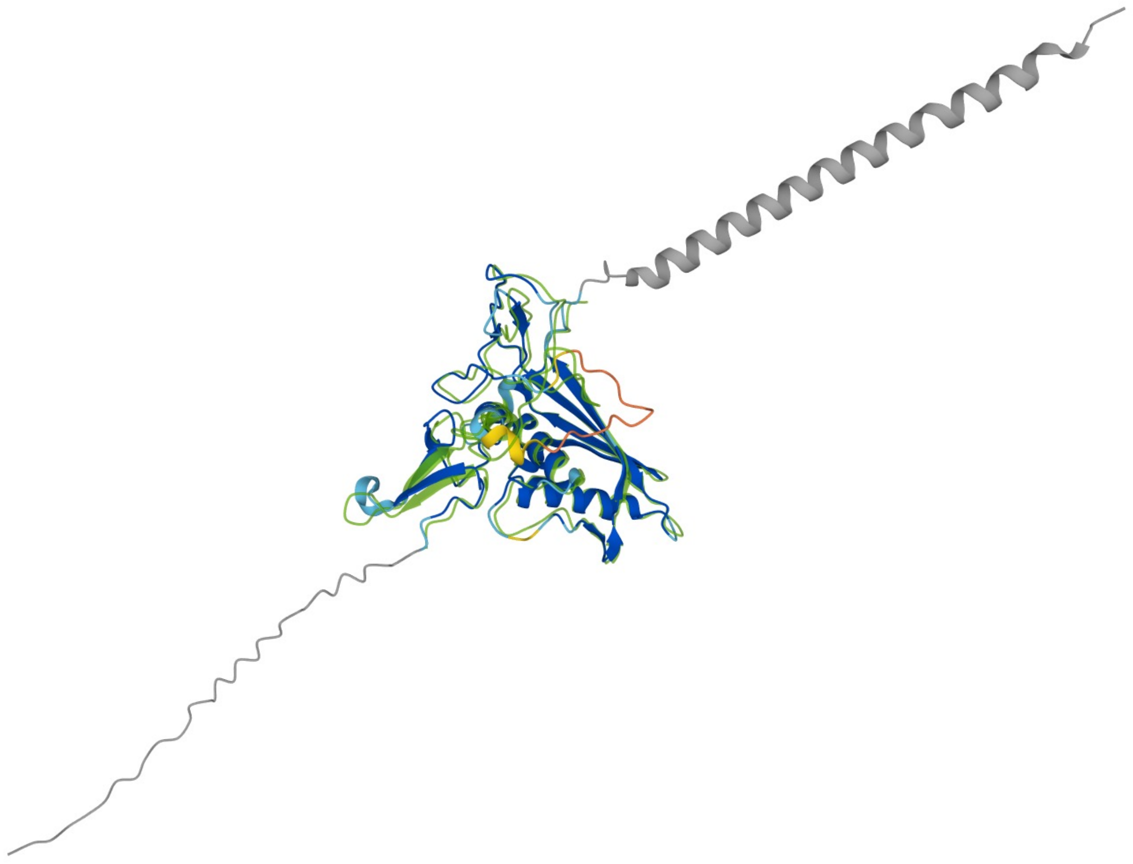
